# The mechanical arthropod vector *Stomoxys calcitrans* influences the outcome of lumpy skin disease virus infection in cattle

**DOI:** 10.1101/2023.03.13.532343

**Authors:** Charlotte G. Cook, Henry Munyanduki, Petra C. Fay, Najith Wijesiriwardana, Katy Moffat, Simon Gubbins, Stuart Armstrong, Carrie Batten, Isabelle Dietrich, David R. Greaves, Karin Darpel, Philippa M. Beard

## Abstract

The poxvirus lumpy skin disease virus (LSDV) is the etiological agent of lumpy skin disease (LSD), a severe disease of cattle and water buffalo that is characterised by numerous necrotic cutaneous nodules. LSD is a rapidly emerging disease, spreading into and across the Middle East, eastern Europe, and Asia in the past decade. The disease causes substantial production and economic losses in rural communities and affected regions. LSDV is mechanically transmitted by haematophagous arthropods including stable flies (*Stomoxys calcitrans*), however our understanding of this mechanical transmission method is sparse. A secreted saliva collection methodology using a modified artificial membrane feeding system was optimised for *S. calcitrans* and used to collect and characterise secreted *S. calcitrans* saliva. Saliva was mixed with LSDV and shown not to affect virus growth in primary bovine fibroblasts. *S. calcitrans* saliva or spot-feeding by *S. calcitrans* was then incorporated into a bovine in vivo experimental model of LSD to determine if either influenced disease pathogenesis. *S. calcitrans* saliva resulted in fewer animals developing disease, however this difference was not statistically significant. Spot-feeding with *S. calcitrans* prior to inoculation did not alter the number of animals that developed disease or the overall severity of disease however disease progression was accelerated as demonstrated by the appearance of cutaneous nodules, detection of viral DNA in the blood stream, and production of neutralising antibodies. This shows that *S. calcitrans* influence disease kinetics through co-incident bite trauma and/or saliva deposition. This increases our understanding of LSDV pathogenesis and highlights the overlooked importance of mechanical vectors in pathogen transmission.

**Author summary:** Insect vectors are important conduits for the transmission of pathogens that cause diseases such as Zika, dengue, malaria, and lumpy skin disease. Biological vector-borne transmission incorporates a replication phase for the pathogen in the insect, whereas no replication occurs in the vector during mechanical transmission. When the insect bites the host it inoculates a pathogen whilst also delivering arthropod-derived factors such as saliva components and causing tissue trauma through biting and probing. Arthropod saliva and/or bite trauma have been shown to enhance the speed and severity of disease following inoculation with a range of biologically transmitted viruses. This study examined if this was true also for the mechanically transmitted pathogen lumpy skin disease virus (LSDV). LSDV is a neglected pathogen that causes severe systemic disease in cattle and is transmitted mechanically by the stable fly *Stomoxys calcitrans*. Using an experimental bovine model of LSD, we found that disease occurred more rapidly when virus was delivered in association with the bites of uninfected flies. This work has increased our knowledge of lumpy skin disease virus transmission, and the discovery that disease outcome can be impacted by previously overlooked mechanical insect vectors should prompt further investigation into this mechanism of transmission.

## Introduction

Arthropod-borne diseases such as dengue, Zika, leishmaniasis, malaria, bluetongue, Rift Valley fever and lumpy skin disease exert a major burden on human and animal health. A number of these arthropod-borne diseases are increasing in incidence and geographical range and moving into temperate zones, representing a growing threat to a warming world (1, 2, 3).

Lumpy skin disease (LSD) is an emerging, high consequence disease of cattle and water buffalo. The causative agent is lumpy skin disease virus (LSDV), genus *capripoxvirus*, family *Poxviridae*. LSD is characterised by multifocal necrotic skin lesions, fever, weight loss, milk drop and death. Morbidity is approximately 10% and mortality 1% (4, 5, 6, 7, 8). LSD contributes to food insecurity and rural poverty and causes substantial economic loss to affected regions through loss of food and draught animal power, lost production, cost of control measures, and trade restrictions (9, 10, 11). LSD was first described in cattle populations in southern Africa in the early 20^th^ century, from where it has spread steadily northwards into northern Africa and the Middle East (12). In 2013 the disease began a rapid period of spread known as the Eurasian LSD epidemic, spreading through cattle populations in Europe, Russia and Asia. Despite its swift emergence and broad impact LSDV remains a neglected pathogen, with fundamental knowledge gaps preventing the design and implementation of effective control programmes.

LSDV is transmitted by haematophagous arthropod vectors. A number of other poxviruses are also transmitted by blood-feeding vectors including the leporipoxviruses myxoma virus (13, 14) and Shope fibroma virus (15, 16), the avipoxvirus fowlpox virus (17, 18), and the capripoxviruses goatpox virus and sheeppox virus (19, 20). LSDV has been transmitted from a clinical to naïve animal under experimental conditions by *Aedes aegypti* mosquitoes (21, 22), *Amblyomma hebraeum* and *Rhipicephalus appendiculatus* ticks (23, 24), and *S. calcitrans* flies (22, 25, 26). LSDV can be acquired from cutaneous nodules by a range of insect species, then retained on the vector mouthparts for days before being deposited in a recipient animal (27, 28). Poxviruses, including LSDV, are believed to be transmitted mechanically with no viral replication taking place in the arthropod vector. This is in contrast to biologically transmitted pathogens which undergo replication in the vector.

The bite of an arthropod vector inoculates a pathogen into the host whilst also delivering arthropod-derived factors such as saliva components, and causing tissue trauma through biting and probing. Arthropod saliva and/or bite trauma have been shown to enhance the kinetics and severity of disease caused by a range of biologically-transmitted pathogens including West Nile virus (29, 30, 31), dengue virus (32, 33), bunyamwera virus (34), Semliki Forest virus (34, 35) and leishmania (36). The mechanisms underpinning arthropod disease enhancement include the *Aedes aegypti* venom allergen-1 which enhances viral replication in macrophages and dendritic cells (37), and *Ae. aegypti* protein sialokinin which increases blood vessel permeability and availability of virus permissive cells (38). While arthropod enhancement of disease has been shown for a number of biologically-transmitted pathogens, this potentiation has not been described for disease caused by mechanical vectors.

*S. calcitrans*, also known as stable flies, are telmophagous filth flies and a common arthropod pest particularly of cattle. They are pool feeders with coarse labellar teeth that create a haematoma in the skin from which they feed (39). Stable flies are found across the world (40) and are most active from spring to mid-autumn with populations peaking in summer (41, 42, 43, 44). *S. calcitrans* tend to remain in areas with a high concentration of cattle, however research has shown that they have the capacity for long range flight up to 225 km (45). *S. calcitrans* feed quickly, engorging after 3-4 minutes although feeds are often interrupted (46). Their interrupted feeding and refeeding behaviour make *S. calcitrans* ideal vectors for pathogen transmission. They are known to be involved in the transmission of equine infectious anaemia virus (47), West Nile virus (48) and trypanosomes (49). They are also suspected to be carriers of Rift Valley fever virus (50) and bovine herpes virus (51). *S. calcitrans* have also been proposed as vectors of LSDV (22, 25, 26, 27, 52, 53).

The interactions of LSDV and the bovine skin in the context of an arthropod bite are unexplored. We hypothesised that, in an analogous fashion to biologically transmitted arthropod pathogens, the saliva and/or bite trauma of a *S. calcitrans* fly would enhance the development of LSD after inoculation of virus into the skin of an animal.

## Results

### Collection and characterisation of *S. calcitrans* secreted saliva

Previous research has described studies of *S. calcitrans* salivary gland extract (SGE) (54, 55, 56). SGE often contains non-salivary structural contaminants therefore we developed a method for collecting secreted *S. calcitrans* saliva to minimise contamination. The method adapted a protocol for collecting secreted saliva from *Culicoides* midges using a heated membrane feeding system (Hemotek™) incorporating a polyvinylidene fluoride (PVDF) membrane (57, 58). A similar artificial membrane feeding system containing 1mM ATP/endotoxin free water solution was used to collect *S. calcitrans* secreted saliva (**Figure 1**). A pot of 25 flies was fed on the PVDF membrane within the feeding system, with each feeding session lasting 10-15 min. Each PVDF membrane was fed on by 3-4 pots of flies sequentially before being retrieved and stored at 4^°^C in PBS. Up to 144 PVDF membranes were fed on during each saliva collection procedure. At the end of the procedure the secreted saliva was collected from the membranes by washing, and concentrated to between 107-380 µg/mL in a total volume of approximately 1000 µL.

**Figure 1.**
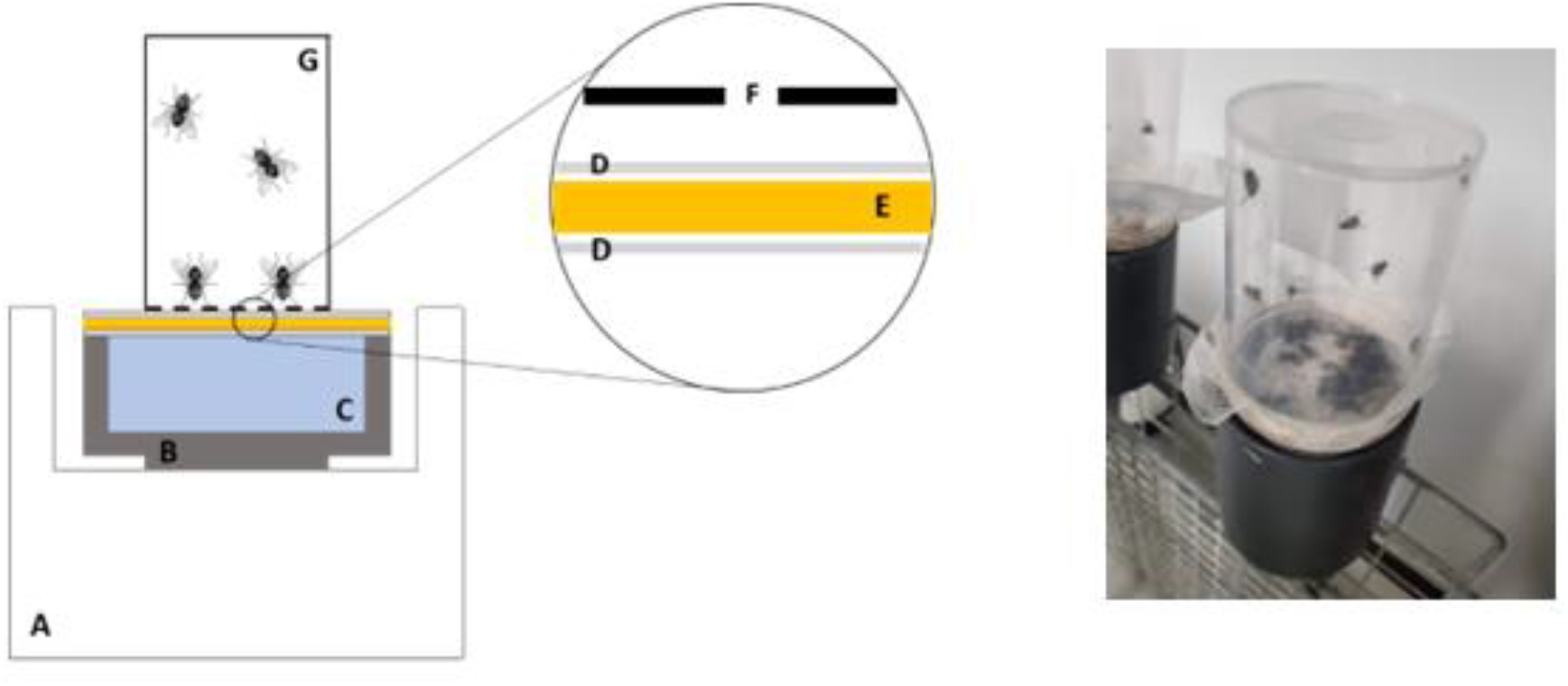
Method for collecting secreted *S. calcitrans* saliva. Left: A: Hemotek™ base heats the reservoir to 37°C; B: screw-in reservoir; C: distilled water to conduct heat to the membrane; D: Parafilm to prevent contamination of the membrane with non-salivary proteins and seal water in the reservoir; E: Durapore PVDF 0.2 μm membrane soaked in ATP solution; F: fine netting to allow flies to feed; G: transparent 4cm diameter pot containing *S. calcitrans* flies. Right: a pot of flies feeding on the apparatus.

Proteins in the concentrated *S. calcitrans* secreted saliva were separated by SDS-PAGE and visualised using silver stain. Six protein bands were consistently visible in each preparation (**Figure 2A**), with the most abundant protein detected at 23-24kDa. Mass spectrometry was used to identify the proteins present within the *S. calcitrans* secreted saliva (**Table 1**). This revealed that the secreted saliva contained types of proteins typical of haematophagous arthropod sialomes, such as endonucleases, serine proteases and carboxylic ester hydrolases (59, 60, 61, 62, 63). The most abundant proteins (determined by count of spectra) were an SCP domain-containing protein and antigen-5. These two proteins have a very high amino acid sequence similarity and may be the same protein. Antigen-5 is a member of the CAP (cysteine-rich secretory proteins, antigen-5 and pathogenesis-related protein 1) superfamily and in the saliva of haematophagous insects has been shown to function as antioxidant, to inhibit collagen-induced platelet aggregation and neutrophil oxidative burst in hosts (64). Salivary antigen-5 was also found to be highly antigenic and to induce antibody-mediated immune responses in repeatedly-exposed hosts (65), and specific immune responses and immunoglobulin binding in cattle (56).

**Figure 2.**
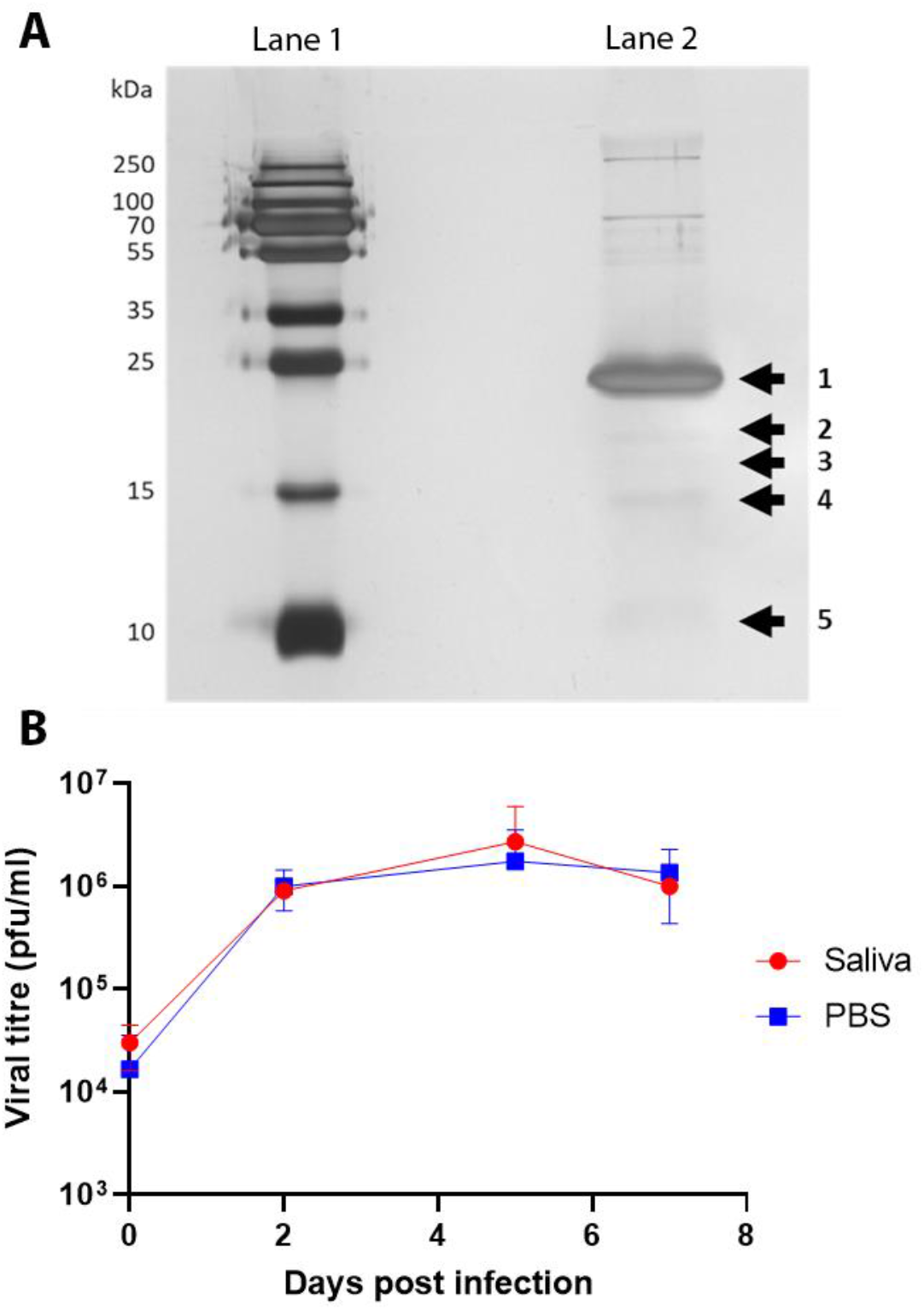
(A) The protein profile of *S. calcitrans* saliva. Secreted saliva from *S. calcitrans* was collected and concentrated then separated on a 15% SDS/PAGE gel before being visualised by silver staining (lane 2). PageRuler Plus was used as the ladder (lane 1). Image is representative of the three collections. (**B). Secreted saliva does not impact LSDV kinetics in BPF.** LSDV was incubated with either secreted *S. calcitrans* saliva or PBS for 2h at RT, then inoculated onto confluent monolayers of BPFs at an MOI of 5. Cells and supernatant were collected together at the time points indicated, sonicated, and titred on MDBK cells. Data shown as mean ± SEM, and are representative of two independent experiments.

**Table 1.** Proteins identified in secreted *S. calcitrans* saliva. Proteins identified in at least two out of three independent preparations were included. Hits were sorted by average abundance determined by spectra counts. Duplicate hits with alternative UniProt accessions are marked in red.

| UniProt Accession | UniProt Annotation | Vectorbase Annotation | Average protein mass (Da) | Average abundance (Spectra) |
| --- | --- | --- | --- | --- |
| A0A1I8PC31 | SCP domain-containing protein |  | 29069 | 443 |
| Q7Z0B5 | Antigen 5 |  | 28940 | 441 |
| A0A1I8Q1A7 | Uncharacterized protein |  | 142581 | 57 |
| A0A1I8PUN2 | Uncharacterized protein | Serine protease 53-like | 79843 | 46 |
| A0A1I8PL64 | Uncharacterized protein | Protein starmaker-like | 55798 | 33 |
| A0A1I8Q6W6 | Alpha-amylase |  | 55421 | 28 |
| A0A1I8PV60 | Uncharacterized protein |  | 45452 | 22 |
| A0A1I8P4Z0 | Uncharacterized protein |  | 35563 | 22 |
| A5WXR8 | Putative 15.6 kDa secreted salivary gland protein |  | 17541 | 14 |
| A0A1I8PSA1 | Lipocln_cytosolic_FA-bd_dom domain-containing protein |  | 21441 | 12 |
| A0A1I8PU36 | Peptidase S1 domain-containing protein |  | 26684 | 12 |
| A0A1I8PR12 | Transferrin |  | 71124 | 11 |
| A0A1I8Q544 | Peptidase S1 domain-containing protein |  | 27388 | 11 |
| A5WXR7 | Putative 13.7 kDa secreted salivary gland protein |  | 15838 | 9 |
| A0A1I8NWI4 | Uncharacterized protein |  | 18501 | 9 |
| A0A1I8P7C1 | Carboxylic ester hydrolase |  | 64867 | 8 |
| A0A1I8Q0V7 | Lipase domain-containing protein |  | 42932 | 8 |
| A0A1I8PQD6 | Uncharacterized protein | Murinoglobulin-2-like | 165805 | 8 |
| A0A1I8PQI3 | Uncharacterized protein | Murinoglobulin-2-like | 166711 | 8 |
| A0A1I8PU39 | Uncharacterized protein | Actin-5C | 41822 | 8 |
| A0A1I8NR78 | Uncharacterized protein | Transmembrane protease serine 9 | 52998 | 6 |
| A0A1I8PGD9 | Uncharacterized protein | Venom serine protease | 44853 | 6 |
| A0A1I8Q114 | Carboxylic ester hydrolase |  | 64782 | 6 |
| A0A1I8PI66 | Uncharacterized protein |  | 13078 | 5 |
| A0A1I8Q1H9 | P/Homo B domain-containing protein |  | 172722 | 5 |
| A0A1I8Q1I4 | P/Homo B domain-containing protein |  | 177770 | 5 |
| A0A1I8QE72 | Endonuclease_NS domain-containing protein |  | 43497 | 5 |
| A0A1I8NTR2 | Peptidase S1 domain-containing protein |  | 27598 | 5 |
| A0A1I8NQD5 | C-type lectin domain-containing protein |  | 21272 | 4 |
| A0A1I8PZ42 | Uncharacterized protein | Protein yellow | 48527 | 4 |
| A0A1I8PZ57 | Uncharacterized protein | Protein yellow | 40878 | 4 |
| A0A1I8PLH0 | Uncharacterized protein | Glycerophosphocholine phosphodiesterase GPCPD1 | 77514 | 4 |
| A0A1I8PLL7 | Uncharacterized protein | Glycerophosphocholine phosphodiesterase GPCPD1 | 75000 | 4 |
| A0A1I8PLN8 | Uncharacterized protein | Glycerophosphocholine phosphodiesterase GPCPD1 | 77642 | 4 |
| A0A1I8PLP5 | Uncharacterized protein | Glycerophosphocholine phosphodiesterase GPCPD1 | 74872 | 4 |
| A0A1I8PQF4 | Uncharacterized protein | Murinoglobulin-2-like | 166546 | 4 |
| A0A1I8PQL1 | Uncharacterized protein | Murinoglobulin-2-like | 165871 | 4 |
| A0A1I8PUP6 | C-type lectin domain-containing protein |  | 19344 | 4 |
| A0A1I8P599 | Peptidase S1 domain-containing protein |  | 28478 | 3 |
| A0A1I8Q0V1 | Uncharacterized protein | Tyrosine-protein phosphatase non-receptor type 23 | 39376 | 3 |
| A0A1I8Q5Q1 | Uncharacterized protein | Protein 5NUC-like | 63674 | 3 |
| A0A1I8P1G4 | Peptidase S1 domain-containing protein |  | 27229 | 2 |

To further characterise the composition of the *S. calcitrans* sialome functional enrichment analysis was performed (Table 2). In accordance with the biological function of saliva, overrepresented molecular functions were serine-type peptidase activity-related functions. “Extracellular region” was overrepresented as cellular component, indicating that the dataset contained several secreted proteins. Once characterised, the *S. calcitrans* secreted saliva was taken forward to characterise the influence of *S. calcitrans* on LSDV *in vitro* and *in vivo*.

**Table 2.** Functional enrichment analysis of proteins identified in *S. calcitrans* saliva. UniProt accession numbers listed in Table 1 were queried against the *S. calcitrans* genome using the g:GOSt on the g:Profiler server. Alternative accession numbers for the same protein were automatically removed prior to analysis (highlighted in red in Table 1). The g:SCS algorithm in g:GOSt was selected for multiple testing correction of p-values with an experiment-wide p-value of 0.05 being considered statistically significant.

| GO:MF |  |  |  |  |
| --- | --- | --- | --- | --- |
| Term Name | Term ID | p (adj) | Enrichment score [-log<sub>10</sub> p (adj)] | Sialome proteins |
| serine-type endopeptidase activity | GO:0004252 | 4.18E-05 | 4.379 | 9 |
| serine-type peptidase activity | GO:0008236 | 8.32E-05 | 4.080 | 9 |
| serine hydrolase activity | GO:0017171 | 8.32E-05 | 4.080 | 9 |
| endopeptidase activity | GO:0004175 | 0.004 | 2.388 | 9 |
| hydrolase activity | GO:0016787 | 0.014 | 1.861 | 17 |
| GO:CC |  |  |  |  |
| Term Name | Term ID | p (adj) | Enrichment score | Sialome proteins |
| extracellular region | GO:0005576 | 0.011 | 1.966 | 3 |

### *S. calcitrans* saliva does not influence the replication of LSDV *in vitro*

Arthropod saliva has been shown to influence the infectivity of viruses in mammalian cell culture, for example *Culicoides* saliva proteins reduced BTV-1 infectivity 2-6 fold in BHK cells but increased infectivity in KC cells (66), Vesicular stomatitis virus growth in *Ae. triseriatus* SGE-treated mouse fibroblast cells was significantly increased compared to untreated controls (67), and crude mosquito saliva inhibited DENV infection of human dendritic cells (68). Therefore, the impact of *S. calcitrans* saliva on LSDV viability *in vitro* was assessed using a one-step growth curve on bovine primary fibroblasts. Varying concentrations of secreted *S. calcitrans* saliva or PBS were combined with LSDV, incubated at room temperature (RT) for 2 h, and then used to infect bovine primary fibroblasts at a MOI of 5. As shown in **Figure 2B**, the addition of *S. calcitrans* saliva to LSDV did not result in any difference in the replication of LSDV when compared to the PBS control. This indicates that *S. calcitrans* saliva does not influence the kinetics or magnitude of LSDV replication in vitro. The influence of *S. calcitrans* in vivo was then studied.

### Spot-feeding by *S. calcitrans* prior to viral inoculation influences the kinetics of LSD

An experimental bovine model of LSD was used to investigate whether *S. calcitrans* saliva or bite trauma influences the infectivity, kinetics, or severity of LSD. Thirty male Holstein-Friesian calves were randomised into three groups of 10 then challenged with LSDV using three different inoculation routes. Group 1 were inoculated with 1×10^6^ pfu of LSDV inoculated intradermally into the skin via 20 “microdoses” of 100 µL. Ten microdoses were injected into both the left and right dorsal flanks of the calf just caudal to the scapula. These 100 µL “microdoses” were designed to imitate inoculation of virus by *S. calcitrans*. Group 2 were inoculated with 1×10^6^ pfu of LSDV mixed with 10 µg of *S. calcitrans* secreted salivary protein, delivered in a total volume of 2 mL in the same 20 x 100 µL intradermal microdose format (**Figure 3A**). Group 3 were spot-fed with approximately 25 adult virus-negative, laboratory reared *S. calcitrans* flies on the left and right craniodorsal flanks of the calves for 5 min (**Figure 3B**), before 1x10^6^ pfu LSDV was inoculated intradermally in 10 microdoses into each feeding site. Examination of the flies immediately post-feeding revealed reddened abdomens (**Figure 3C**), consistent with biting and intake of blood. Calves in all three groups were monitored and samples collected for 28 days after inoculation.

**Figure 3.**
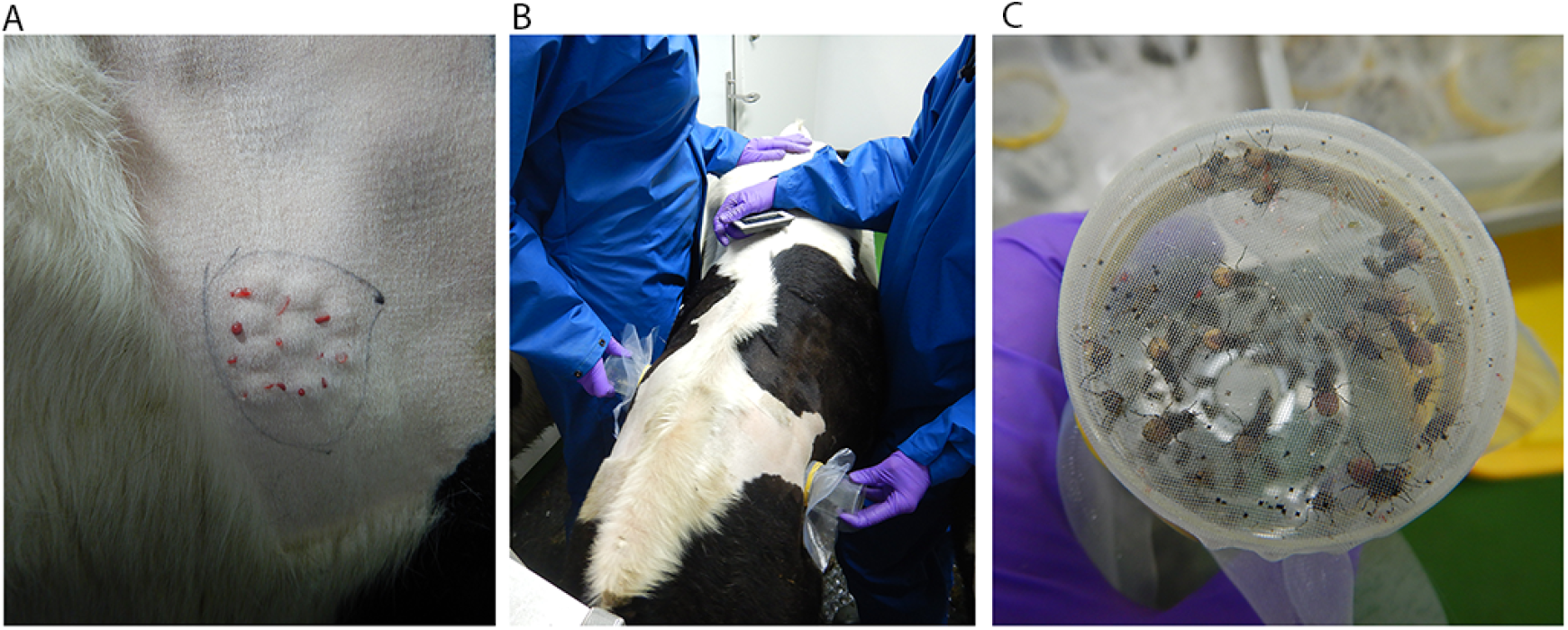
LSDV inoculation methodology. Cattle were inoculated using three different methods. (A) Cattle in group 1 were inoculated intradermally with 1 x10^6^ pfu/ml LSDV in 10 microdoses of 100µl on the left and right craniodorsal flank, just caudal to the scapula. Cattle in group 2 were inoculated intradermally in the same manner in 10 microdoses of 100µl, but with 1x10^6^ pfu/ml LSDV mixed with 10µg *S. calcitrans* salivary proteins. The image shows microblebs from an inoculation site of a calf in group 2, with the site outlinned in black ink. (B) Cattle in group 3 were fed on by pots of *S. calcitrans* flies on the left and right craniodorsal flank, as shown in the image, then inoculated intradermally with 1 x10^6^ pfu/ml LSDV in 10 microdoses of 100µl (the same as group 1 and 2) into the sites of feeding. (C) Post-feeding the flies that fed on the cattle in group 3 showed evidence of feeding (red abdomens).

Clinical disease, defined as the development of one or more cutaneous nodules distant from the site of inoculation, occurred in 16 of the 30 calves. Six out of ten animals inoculated with virus alone (group 1), three out of ten animals inoculated with a mixture of virus and *S. calcitrans* saliva (group 2), and seven out of ten animals spot-fed by flies then inoculated with virus (group 3) developed clinical disease (**Figure 4**). These frequencies do not differ between the groups (Fisher exact test: P=0.27). The gross pathology was similar in all three groups with firm, slightly raised cutaneous lesions first detected 5-7 dpi and developing from papule to vesicle to nodule before becoming necrotic by approximately 12 dpi. This was consistent with previous bovine experimental models of LSD (69). Severely affected calves developed over 100 cutaneous nodules. The clinical calves also developed lymphadenopathy and pyrexia. Severely affected calves were euthanased between 9-13 dpi.

**Figure 4.**
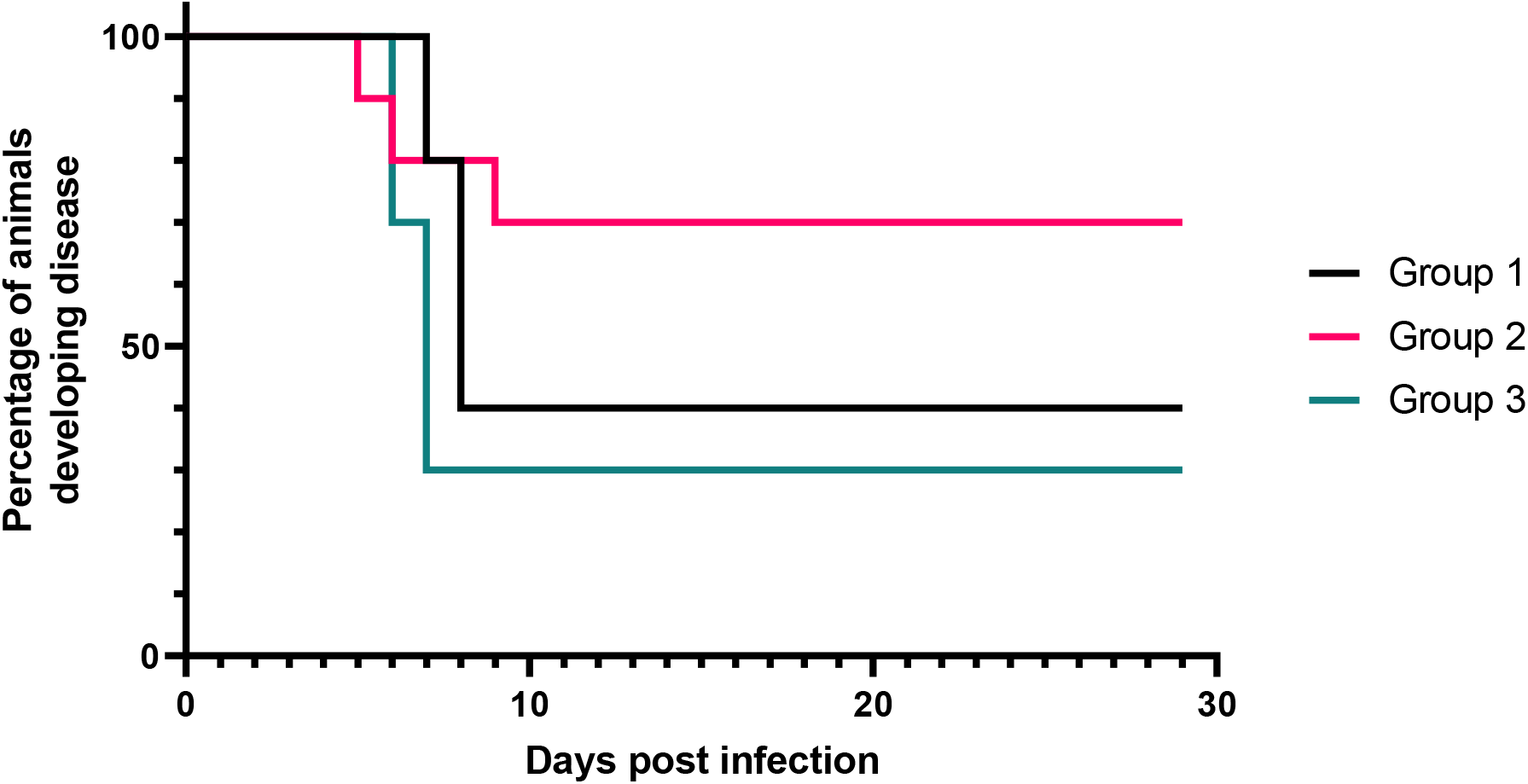
Clinical outcome of different inoculation strategies of LSDV. Disease development was tracked in each animal by the initial onset of cutaneous nodule development and shown as a Kaplan-Meier graph. Those that developed nodules were scored as “1” and those that did not were scored as “0”. Data were tested for significant differences between the groups using the log-rank test, but were not significant (P=0.154).

Fourteen of the 30 calves inoculated with LSDV did not develop any cutaneous nodules distant from the inoculation site and were classified as non-clinical. These calves did exhibit an increase in size of pre-scapular and pre-femoral lymph nodes from 3-15 dpi and, briefly, a raised temperature around 5-9 dpi. Overall, this study shows that the “microdose” method of inoculating LSDV produces lumpy skin disease in 60-70% of challenged calves.

We then examined whether the addition of *S. calcitrans* saliva or spot feeding by *S. calcitrans* altered the infectivity of LSDV or the course of disease. The outcome of inoculation – disease or no disease – differed between the groups as described above, with only 3 calves in group 2 developing clinical disease, compared to 6 in group 1 and 7 in group 3. However this trend did not reach statistical significance (using the log-rank test, *P*=0.154, **Figure 4**). No significant differences were noted between the three groups when the temperature, lymphadenopathy, heart rate, respiration rate, and demeanour were compared (data not shown). However significantly more cutaneous nodules per clinical animal in group 3 were present at 7 dpi compared to group 1 (mixed effects model, *P*=0.013, **Figure 5**), indicating that spot-feeding ahead of virus inoculation accelerates the appearance of disease. This accelerated disease was also evident in a clinical scoring system that takes into account the key signs of disease (**Table 3**). The clinical animals in group 3 reached a high severity score more quickly than groups 1 and 2, and significantly higher than group 1 at 7dpi (*P*=0.010, mixed effects model, **Figure 6**). The clinical calves in group 2 had a lower disease severity over the course of the experiment, but the variation between the three clinical cattle in this group prevented any statistical significance from being reached. Overall, these results indicate that fly feeding prior to viral inoculation accelerates the kinetics of LSD.

**Figure 5.**
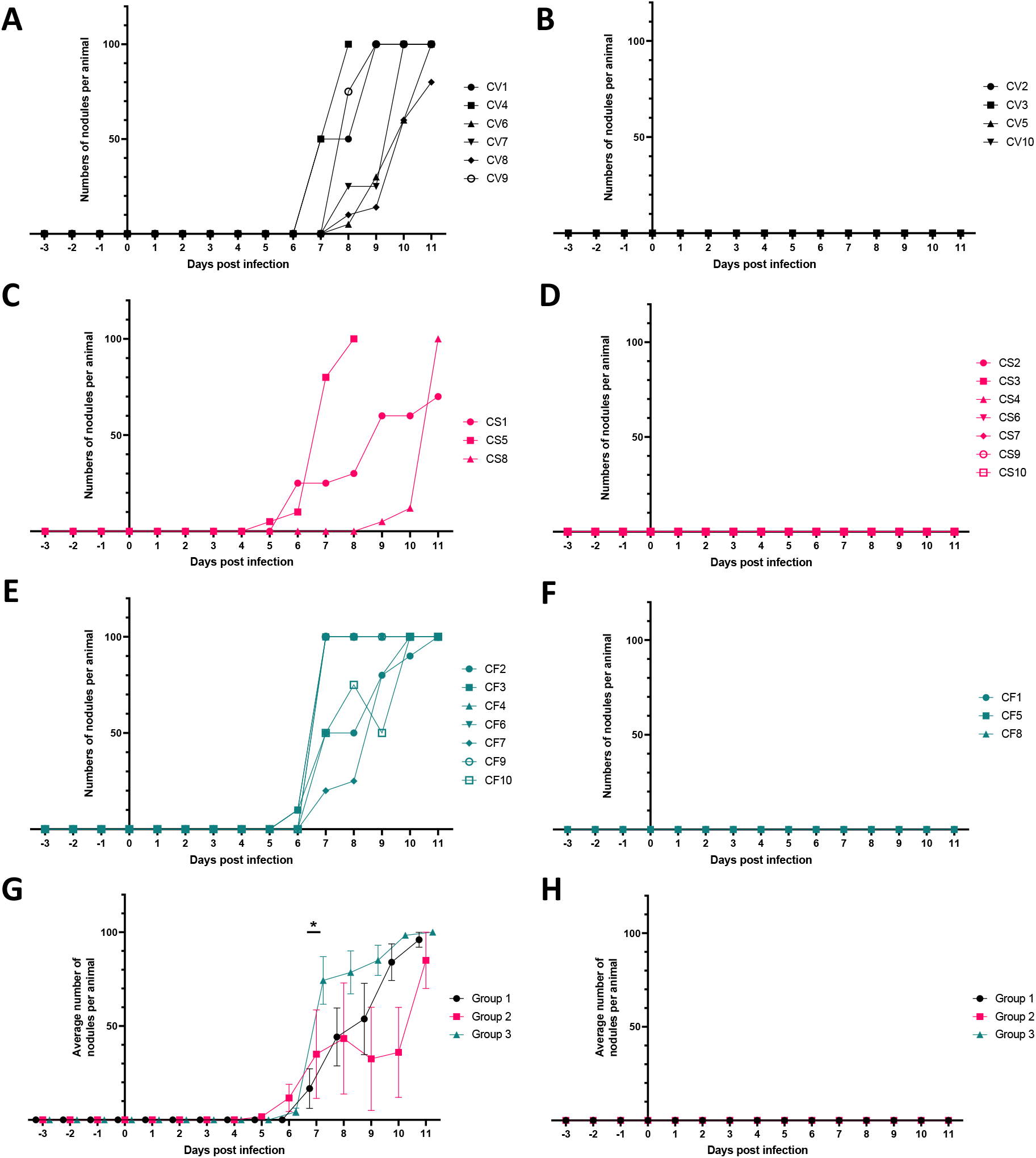
Cutaneous nodules appear faster in the calves in group 3 that were spot-fed before virus inoculation. Nodules were measured by manual examination of each animal, up to a maximum of 100 nodules per animal. Calves with cutaneous nodules distant from the site of inoculation were designated clinical (left column) and calves that did not develop cutaneous nodules distant from the site of inoculation were designated nonclinical (right column). (A and B) number of nodules on each calf from group 1 (inoculated with LSDV alone), (C and D) number of nodules on each calf from group 2 (inoculated with LSDV and secreted *S. calcitrans* saliva), (E and F) number of nodules on each calf from group 3 (inoculated with LSDV into area that had been previous spotfed by *S. calcitrans* flies). (G and H) Average number of nodules per animal (black for group 1, pink for group 2, and blue group 3). Data shown as mean ± SEM. Graphs show data collected from -3dpi to 11dpi. Significance was determined by a mixed effects model.

**Table 3.**
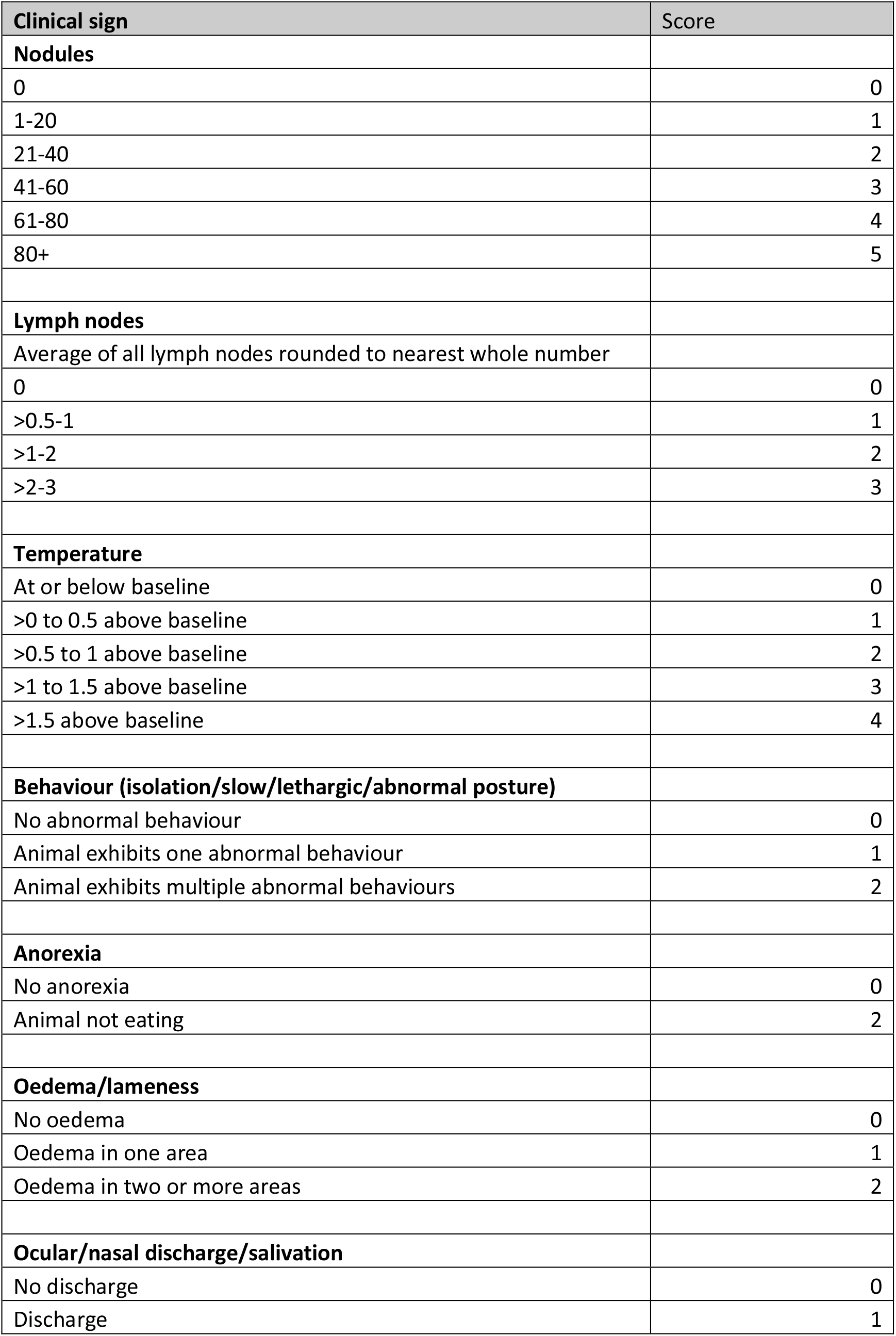
Clinical scoring system for lumpy skin disease

| Clinical sign | Score |
| --- | --- |
| **Nodules** |  |
| 0 | 0 |
| 1-20 | 1 |
| 21-40 | 2 |
| 41-60 | 3 |
| 61-80 | 4 |
| 80+ | 5 |
| **Lymph nodes** |  |
| Average of all lymph nodes rounded to nearest whole number |  |
| 0 | 0 |
| >0.5-1 | 1 |
| >1-2 | 2 |
| >2-3 | 3 |
| **Temperature** |  |
| At or below baseline | 0 |
| >0 to 0.5 above baseline | 1 |
| >0.5 to 1 above baseline | 2 |
| >1 to 1.5 above baseline | 3 |
| >1.5 above baseline | 4 |
| **Behaviour (isolation/slow/lethargic/abnormal posture)** |  |
| No abnormal behaviour | 0 |
| Animal exhibits one abnormal behaviour | 1 |
| Animal exhibits multiple abnormal behaviours | 2 |
| **Anorexia** |  |
| No anorexia | 0 |
| Animal not eating | 2 |
| **Oedema/lameness** |  |
| No oedema | 0 |
| Oedema in one area | 1 |
| Oedema in two or more areas | 2 |
| **Ocular/nasal discharge/salivation** |  |
| No discharge | 0 |
| Discharge | 1 |

**Figure 6.**
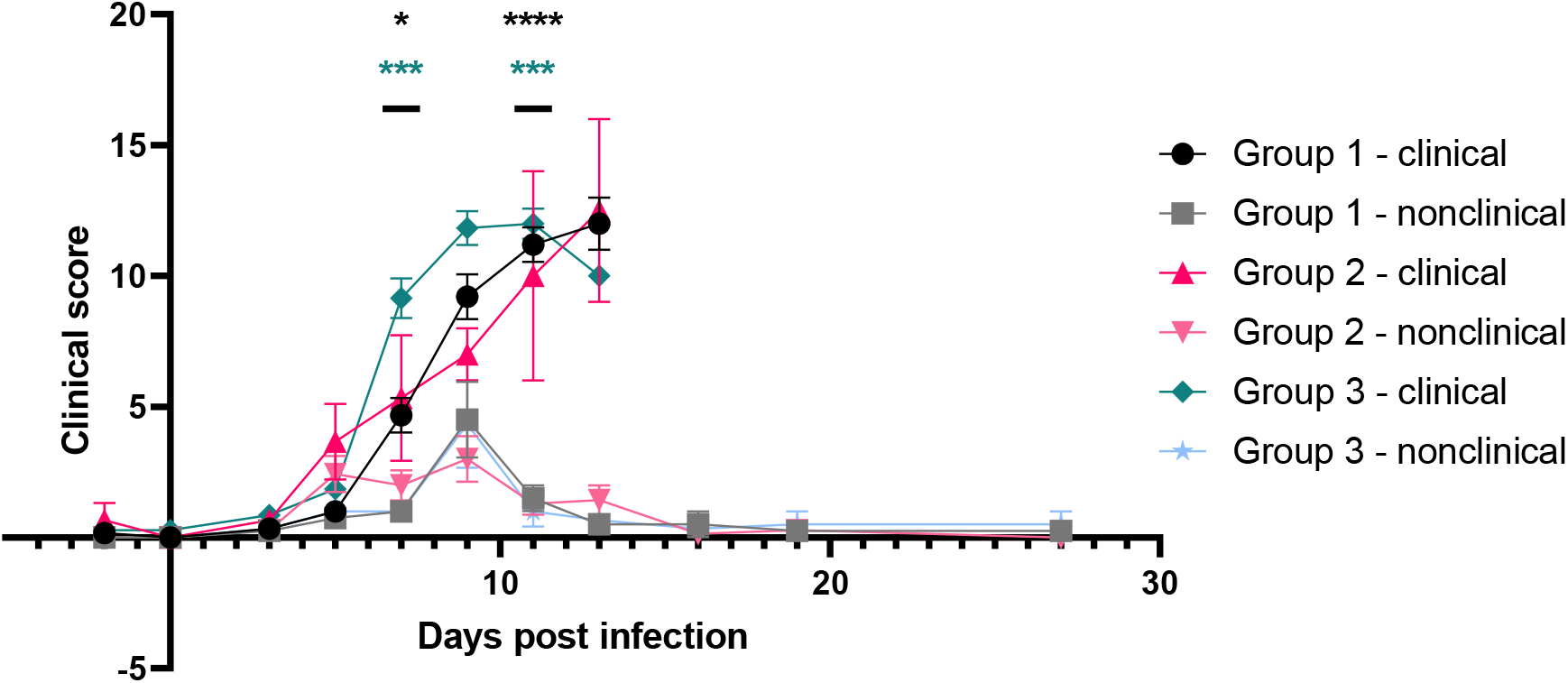
Spot feeding prior to LSDV inoculation results in more rapid development of disease. Clinical scores for each animal were separated according to group (1-3) and into clinical or nonclinical outcomes and averaged. Data shown as mean ± SEM. Data are shown to 13 dpi for clinical animals and 27 dpi for non-clinical animals. The score of the group 3 clinical animals was significantly higher than the score of the group 1 clinical animal at 7dpi (p=0.0104, mixed effects model). A significant difference was also detected between clinical and non-clinical animals in group 1 (black) and between the clinical and non-clinical animals in group 3 (blue/green) at both 7dpi and 11dpi (mixed effects model).

### Spot feeding results in quicker viraemia and more virus in blood

Fly feeding prior to inoculation of LSDV resulted in cutaneous nodules appearing more rapidly, while inoculation of *S. calcitrans* secreted saliva along with the virus reduced the number of calves that developed disease. To explore if these changes correlated with LSDV replication the presence of virus in the blood and skin was quantified.

The amount of viral DNA in the blood of calves in each treatment group was determined by PCR assay targeting viral gene LSDV074. As expected, all 16 clinical animals developed viraemia with increasing amounts of viral DNA (inferred by decreasing cycle threshold numbers) detected in the blood between 5 and 11 dpi. In contrast, viral DNA was detected in the blood of non-clinical animals only intermittently and at low amounts (**Figure 7** A-C). Median C_T_ values were significantly (P<0.05) lower for clinical compared with non-clinical calves at 9 and 11 dpi (group 1), at 7, 11 and 13 dpi (group 2), and at 7, 9 and 11 dpi for group 3. Statistical analysis of the data revealed that median C_T_ values did not differ significantly (*P*>0.19) between non-clinical calves infected by different methods at any time point.

**Figure 7.**
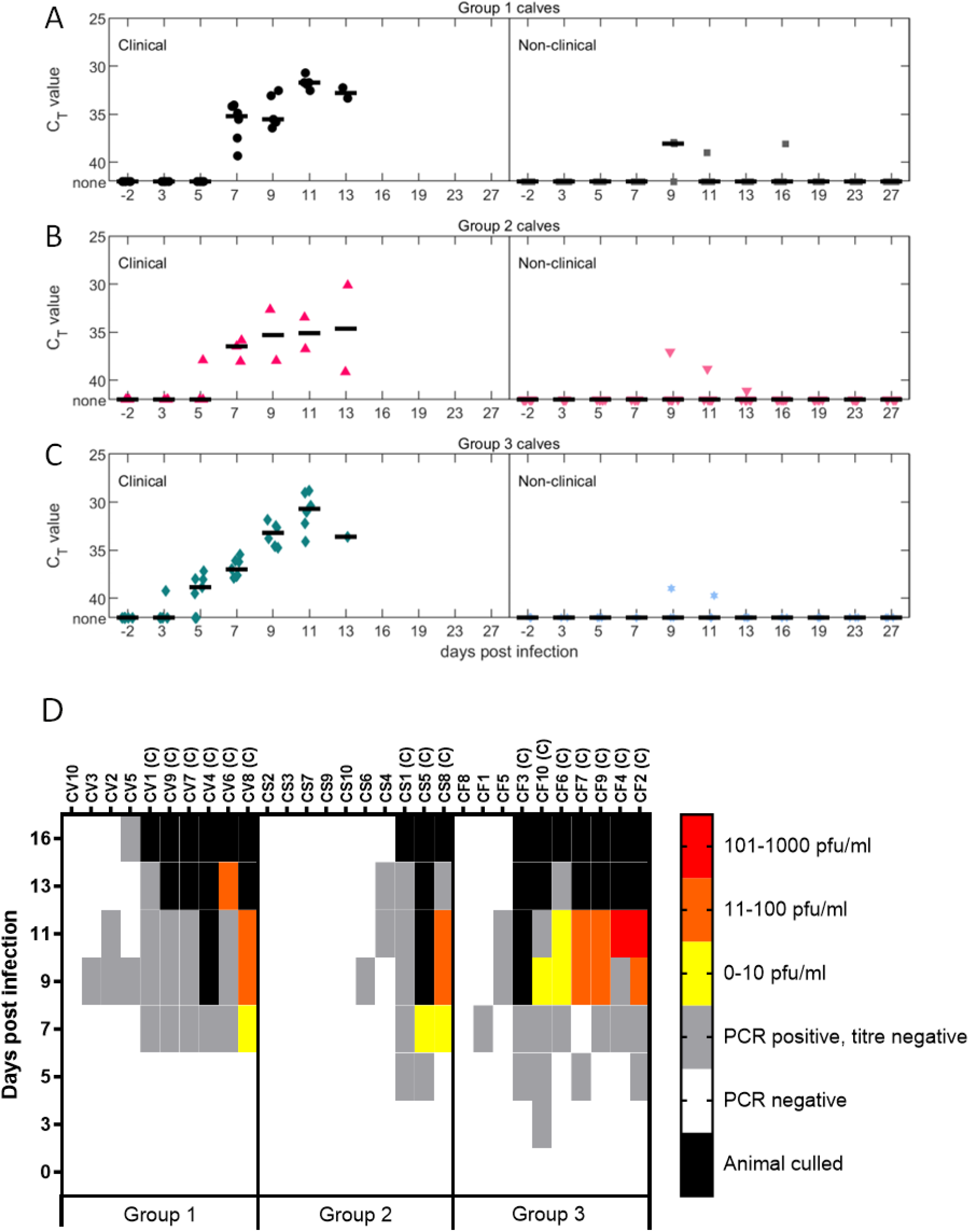
Clinical animals develop a higher viraemia than non-clinical animals. Comparison of C_T_ values in blood at different days post infection in calves in group 1 (A), group 2 (B) and group 3 (C). The left-hand plots show the C_T_ values for clinical calves and the right-hand plots those for non-clinical calves. (D). Blood samples that were positive on PCR were titred on MDBK cells, and viral foci counted manually. A letter C after the animal identification number denotes a clinical animal.

The amount of viral DNA in the blood of clinical calves was compared across the three groups. Viraemia was detected earlier in the calves in group 3 at 3 and 5dpi compared to 5 and 7 dpi for the calves in groups 1 and 2. One calf from group 3 was positive at 3 dpi, and 5 calves from group 3 were positive at 5 dpi. In comparison no calves from group 1 were positive on either 3 or 5 dpi, and only one calf from group 2 was positive at 5 dpi. C_T_ values were significantly (*P*=0.02) lower at 5 dpi for clinical calves from group 3, infected by microdoses of LSDV after feeding by uninfected *S. calcitrans* (median C_T_=38.8) compared with those in group 1 that had been infected by microdoses of LSDV only (median C_T_=42). This reveals that spot feeding with *S. calcitrans* flies accelerates the development of viraemia in LSD.

To quantify the amount of infectious virus present in the blood of the inoculated calves, viral titrations were carried out on the 63 blood samples that contained viral DNA. Of these, 52 samples were from clinical calves and 11 samples from nonclinical calves. Live virus was detected in eighteen samples, all from clinical animals (animals marked (C) in **Figure 7D**). The amount of virus detected was low, ranging from 10-1000 pfu/mL, consistent with previous experimental studies of LSD (69, 70). Live virus was recovered from the blood of two calves from group 1, two calves from group 2, and six calves from group 3. The maximum amount of virus detected in blood samples from groups 1 and 2 was 100 pfu/mL, however higher titres were present in the blood samples from calves in group 3, up to 1000 pfu/mL.

At 5, 11 and 16 dpi a 2mm skin biopsy was taken from each of the calves. The biopsies were taken from normal skin of all calves at 5 dpi, from cutaneous nodules on the clinical animals at 11 and 16 dpi, and from normal skin of nonclinical animals at 11 and 16dpi. DNA was extracted from the biopsy and viral genomic DNA quantified using a qPCR targeting LSDV068. This PCR incorporated a standard curve to enable quantification of the amount of viral DNA present. No viral DNA was detected in any of the skin samples collected at 5dpi, and no viral DNA was detected in any of the biopsy samples taken from the skin of a nonclinical calf at 11 and 16 dpi. In contrast, all samples taken from the nodules of clinical calves at 11 dpi were positive (5/5 clinical animals in group 1, 2/2 clinical animals in group 2, and 6/6 clinical animals in group 3). The viral genome copy numbers present in each microbiopsy at 11 dpi were similar across the inoculation groups (**Figure 8**), with the exception of CV7 which contained a lower amount of viral DNA. Overall, there was no evidence to suggest that the amount of viral DNA in the nodules of clinical animals was impacted by *S. calcitrans* saliva or spot feeding.

**Figure 8.**
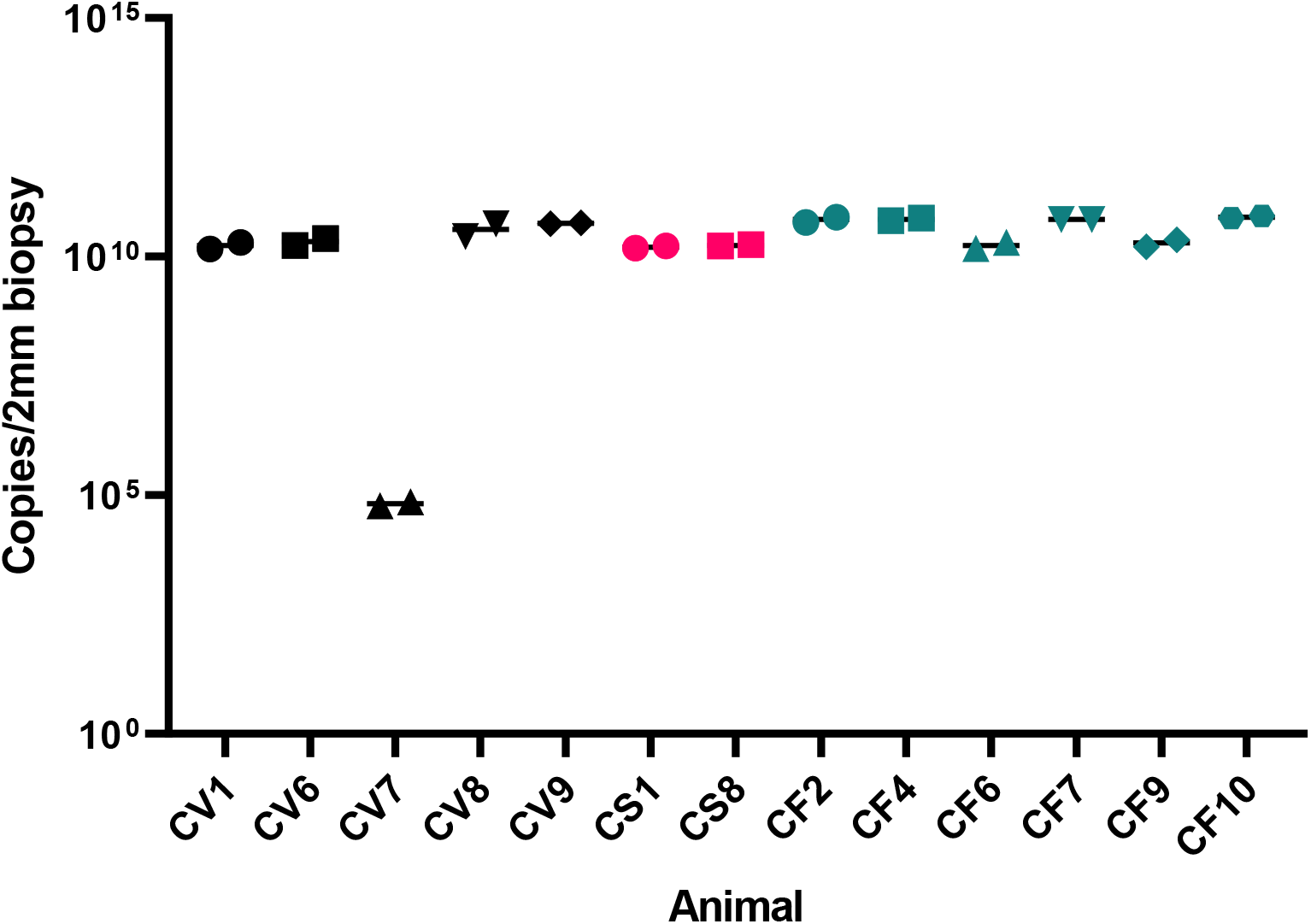
Virus is present in the nodules of clinical animals at 11dpi. Skin samples were taken from using a 2mm biopsy punch at 5dpi, 11dpi and 16dpi. Viral DNA was extracted from the tissue and quantified using qPCR in duplicate. Viral DNA was detected only in 11dpi samples and shown here. Data shown as mean ± SEM.

### *S. calcitrans* saliva and fly feeding does not influence the blood cell population changes in response to LSDV inoculation

The immune response of the challenged calves was studied by examining changes in blood cell populations over time and in the production of neutralising antibodies. The number of mononuclear leukocytes in the venous blood of the calves at -2, 3, 7 and 9 dpi were examined using a flow cytometry panel. The panel incorporated counting beads to enable quantification of absolute numbers of monocytes, T cells, B cells and γδ T cells (**Figure 9**). The cell counts were then presented as a percentage change from the preinfection baseline. Changes in cell numbers were broadly similar across the three groups of calves. A reduction in all four cell subsets was seen at 3 and 7 dpi when compared to -2 dpi. At 9 dpi the number of γδ T cells continued to drop, however the T and B cell numbers increased. At 9 dpi the number of monocytes dropped in group 1, plateaued in group 2, and increased in group 3. There were no statistically significant differences in the number of blood cells between the three groups at any time point.

**Figure 9.**
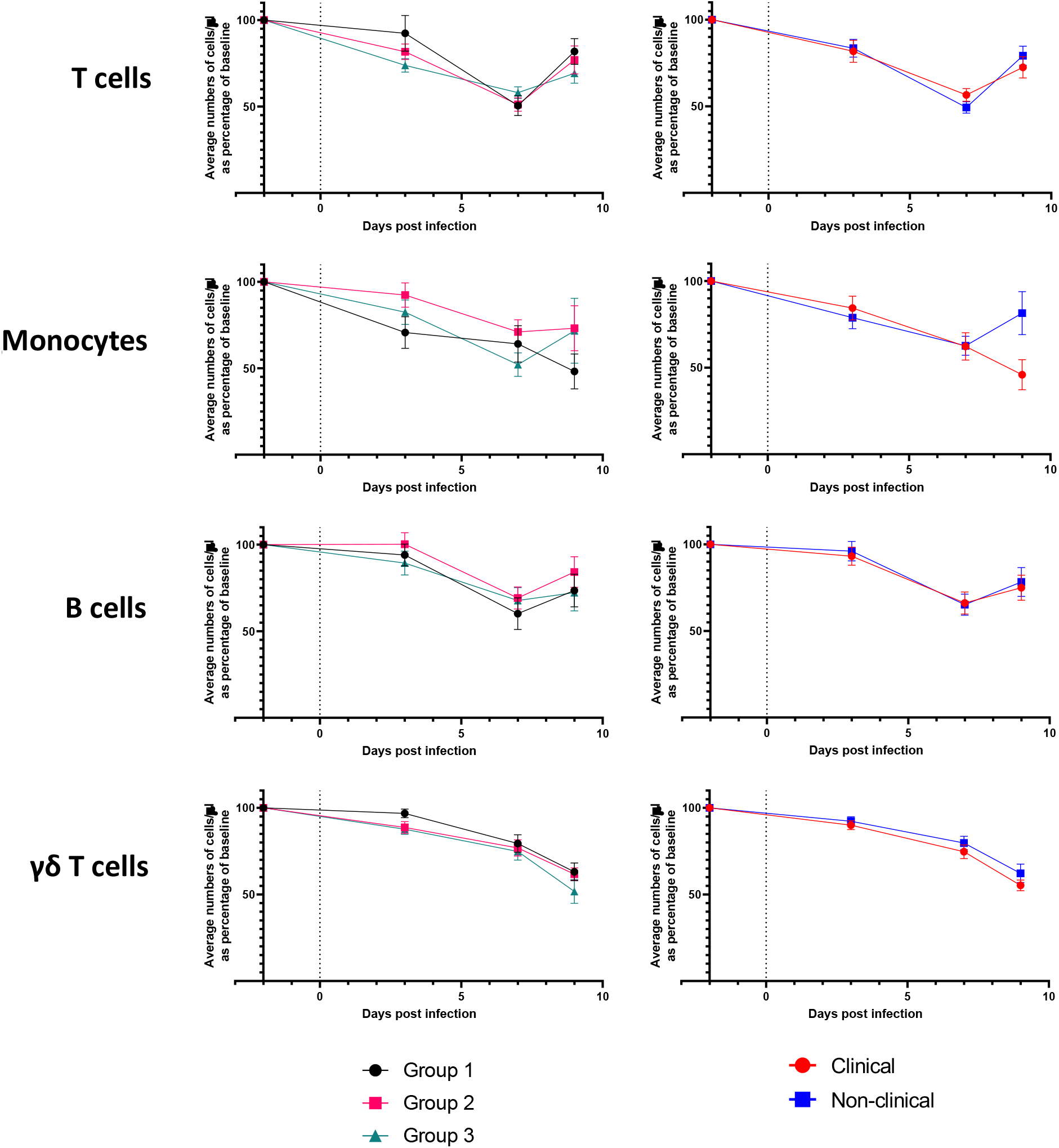
LSDV infection of cattle induces a lymphopenia in clinical and non-clinical animals. For each cell type the triplicate counts of cells/µl for each animal were averaged and converted to a percentage of the cell count at -2 dpi for each animal. These proportions were then pooled into the three groups of calves and averaged (group 1 black, group 2 pink and group 3 green) on the left, and into clinical and non-clinical outcomes on the right. Animals that became clinical are shown in red, and those that remained non-clinical are shown in blue. Data shown as mean ± SEM. The dashed line indicates the day of challenge.

The data were then re-analysed, comparing the clinical and nonclinical calves from all three groups. The absolute numbers of monocytes, T cells, B cells and γδ T cells were remarkably similar in both groups of calves, with the only difference being the number of monocytes at 9 dpi. At 7 dpi the number of monocytes in the clinical calves was 62.3% of the -2 dpi number, and 62.6% of the -2 dpi number in the non-clinical calves. At 9 dpi the number of monocytes in the clinical calves had reduced even further to 45.9% of pre-challenge numbers, but had increased in the non-clinical calves to 81.5%. There were no significant differences (*P*>0.39) in percentage of CD14^+^ cells at any time point with the exception of 9 dpi (*P*=0.009), where the median percentage was lower in clinical (33.8%) compared with non-clinical (62.2%) calves.

### Fly-fed and inoculated animals produced antibodies more rapidly

A fluorescent virus neutralisation test (FVNT) was used to analyse the ability of sera collected at -2, 7, 11, 16 and 27 dpi to completely neutralise LSDV (71). The highest dilution of sera which completely neutralised the virus is shown in **Table 4** with calves that developed clinical disease highlighted in red. The data is plotted in **Figure 10**. There were no significant differences amongst calves infected by different methods at any time point (*P*>0.23), except 11 days post infection (dpi) (*P*=0.007). At this time point (11 dpi) the median titre was significantly (*P*<0.04) higher for group 3 calves that had been infected by microdoses of LSDV following feeding by uninfected *S. calcitrans* (median FVNT=10^1.3^) than for those group 2 calves infected by microdoses of mixed LSDV and secreted *S. calcitrans* saliva (median FVNT=0) or group 1 calves infected by microdoses of LSDV only (median FVNT=0). These data indicate that spot feeding with *S. calcitrans* ahead of LSDV inoculation stimulates a more rapid humoral immune response.

**Figure 10.**
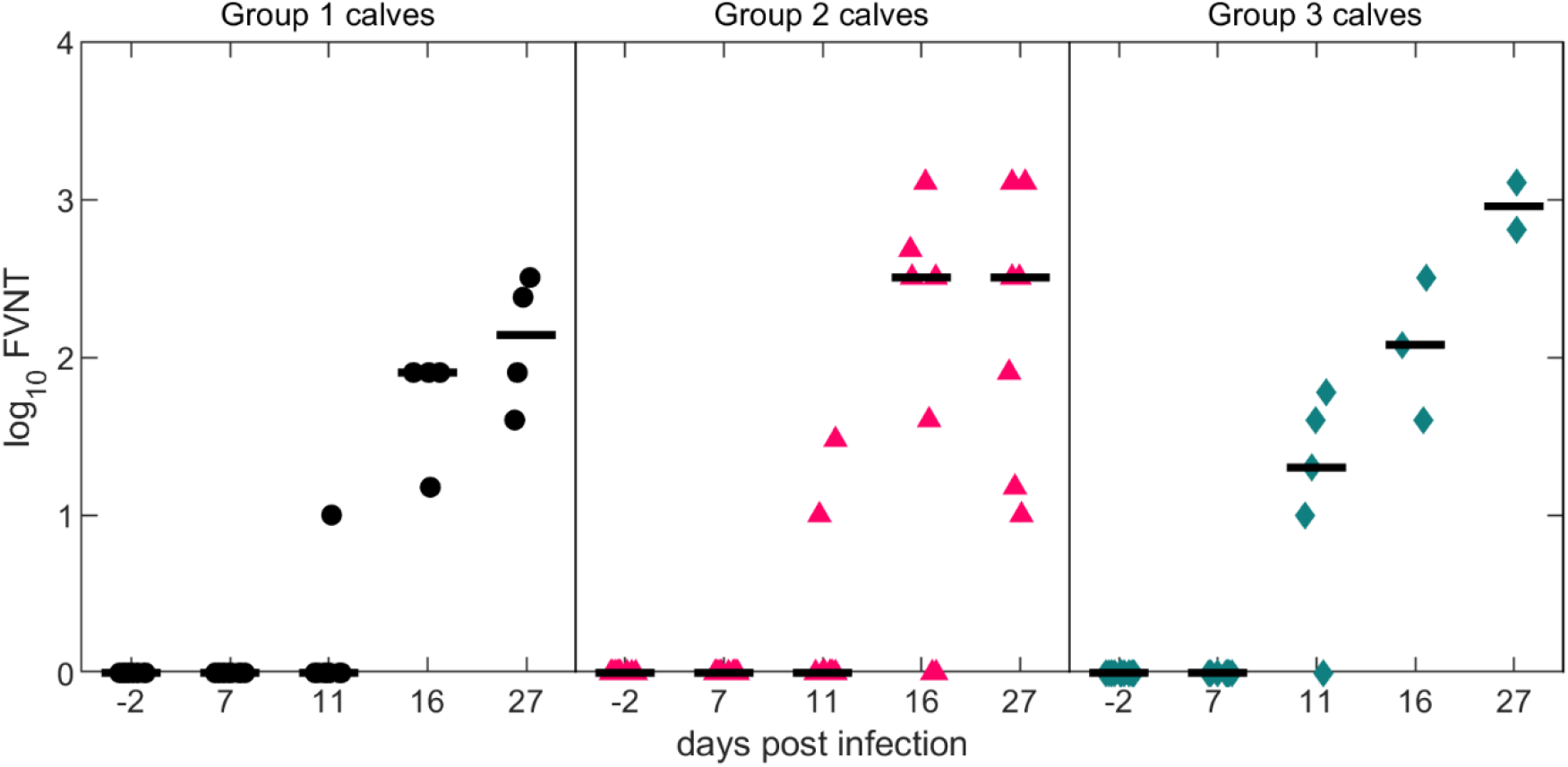
Comparison of log_10_ FVNT over time in calves infected with lumpy skin disease virus (LSDV). Difference colours indicate different infection methods: calves infected via microdoses of LSDV following feeding by uninfected *Stomoxys calcitrans* (group 1, red); calves infected with microdoses of a mixture of LSDV and secreted *S. calcitrans* saliva (group 2, magenta); and calves infected with microdoses of LSDV only (group 3, blue).

**Table 4.** Total neutralisation titres of sera

|  | Complete Neutralisation (FVNT<sub>100</sub>) Days Post Infection |  |  |  |  |
| --- | --- | --- | --- | --- | --- |
|  | -2 | 7 | 11 | 16 | 27 |
| CV1 | neg | neg | neg |  |  |
| CV2 | neg | neg | neg | 15 | 40 |
| CV3 | neg | neg | neg | 80 | 80 |
| CV4 | neg | neg |  |  |  |
| CV5 | neg | neg | neg | 80 | 320 |
| CS1 | neg | neg | neg |  |  |
| CS2 | neg | neg | neg | 320 | 320 |
| CS3 | neg | neg | neg | 40 | 80 |
| CS4 | neg | neg | 10 | 320 | 320 |
| CS5 | neg | neg |  |  |  |
| CF1 | neg | neg | 10 | 40 |  |
| CF2 | neg | neg | 10 |  |  |
| CF3 | neg | neg |  |  |  |
| CF4 | neg | neg | neg |  |  |
| CF5 | neg | neg | 10 | 320 | >1280* |
|  | -2 | 7 | 11 | 16 | 27 |
| CV6 | neg | neg | neg |  |  |
| CV7 | neg | neg | 10 |  |  |
| CV8 | neg | neg | neg |  |  |
| CV9 | neg | neg | neg |  |  |
| CV10 | neg | neg | neg | 80 | 240 |
| CS6 | neg | neg | neg | >1280* | >1280* |
| CS7 | neg | neg | neg | neg | 15 |
| CS8 | neg | neg | 30 |  |  |
| CS9 | neg | neg | neg | neg | 10 |
| CS10 | neg | neg | neg | 480 | >1280* |
| CF6 | neg | neg | neg |  |  |
| CF7 | neg | neg | 60 |  |  |
| CF8 | neg | neg | 10 | 120 | 640 |
| CF9 | neg | neg | 40 |  |  |
| CF10 | neg | neg | 20 |  |  |

## Discussion

Poxviruses including myxoma virus, fowlpox virus and LSDV are transmitted mechanically by a range of blood-feeding arthropod vectors. LSDV is an unusual poxvirus because haematogenous arthropod vectors are its only route of transmission, with direct transmission playing little if any role in the epidemiology of LSD (27, 72, 73, 74). This study investigates the influence of the stable fly *S. calcitrans* on the replication and spread of LSDV in vitro, and the kinetics and severity of LSD in vivo. *S. calcitrans* saliva was found to have no effect on LSDV replication in vitro, however addition of secreted *S. calcitrans* saliva to LSDV inoculum in a bovine experimental model of LSD reduced the proportion of calves that developed clinical disease. In contrast, spot feeding by *S. calcitrans* prior to intradermal viral inoculation *in vivo* accelerated the kinetics of clinical signs of disease, the appearance of viraemia, and the production of neutralising antibodies. This indicates that arthropod-induced enhancement is a factor in disease caused by mechanically-transmitted as well as biologically-transmitted pathogens.

Previous studies of *S. calcitrans* saliva have used salivary gland extract (SGE) which involves dissection of the salivary glands from insects followed by maceration and filtration to minimise cellular debris. SGE of *S. calcitrans* has been used to study the proteome and transcriptome (54) of the salivary gland of the fly and the effect of the components of SGE on the bovine immune response (55, 56). The disadvantage of using SGE is the presence of structural components of the salivary gland which are not usually found in saliva. In contrast, the “secreted” methodology described here minimises contaminants and produces more representative and reproducible *S. calcitrans* saliva. Mass spectrometry of the secreted *S. calcitrans* saliva revealed putative immunomodulatory proteins such as antigen-5 as well as enzymatic components of saliva such as peptidases, endonucleases, and amylases, and furthers our knowledge of the fly saliva. This is consistent with previously published arthropod sialomes. This method for obtaining minimally contaminated *S. calcitrans* saliva will facilitate studies of LSDV and other pathogens that are transmitted by this abundant vector.

We found that the addition of *S. calcitrans* secreted saliva did not influence the replication or spread of LSDV in bovine primary fibroblasts, indicating the saliva did not directly impact the infectivity of the virus, or induce an effective antiviral effect in the bovine fibroblasts. The influence of arthropod saliva on virus replication in cell culture is inconsistent with some studies reporting an impact (66, 67, 68), but others not (38).

Addition of arthropod saliva enhances disease caused by multiple biologically transmitted viruses (29, 30, 31, 32, 36, 38, 67). We hypothesised that adding saliva to a mechanically transmitted pathogen, LSDV, would likewise enhance disease. However, fewer calves inoculated with virus mixed with saliva (group 2) developed clinical disease (3/10) compared to calves in group 1 which received virus alone (6/10), suggesting that *S. calcitrans* saliva inhibits the ability of LSDV to cause disease. Whilst a strong trend, this difference did not reach statistical significance. The amount of saliva added to the inoculum was 0.5 µg of *S. calcitrans* saliva in each microdose, comparable with other studies which have examined the influence of arthropod saliva on disease outcome (29, 38). This may not accurately represent the amount of saliva inoculated during a *S. calcitrans* bite. Likewise the amount of virus in the inoculum, 5 x 10^4^ pfu per microdose, may be greater than the amount inoculated by an arthropod in the field. The dose of LSDV inoculated during an arthropod bite is very small and difficult to quantify because LSDV is a mechanically transmitted pathogen and does not replicate within the arthropod vector. Future studies could examine the impact of varying the amount of *S. calcitrans* saliva or the amount of virus in the inoculum on disease outcome.

Spot-feeding is often used to model arthropod-enhancement of disease as both arthropod saliva inoculation and bite trauma occur (30, 33, 34). *S. calcitrans* is a telmophagous feeder with slashing mouthparts that create a haemorrhagic pool from which to feed (39), and are therefore likely to modulate the skin microenvironment at the bite site. We found that spot feeding prior to LSDV inoculation accelerated the pathogenesis of LSDV in our model of disease, with earlier development of cutaneous nodules, earlier appearance of viral DNA in the blood stream, and earlier production of neutralising antibodies. Characterisation of the changes to the mononuclear blood cell population of the calves in response to LSDV inoculation did not reveal a mechanism to explain the difference seen, with no differences detected between the three groups of calves studied. Possible mechanisms for the influence of spot feeding on the development of LSD are the recruitment of inflammatory cells which promote viral survival, as seen in the recruitment of neutrophils to the skin in response to sand fly bites promoting leishamaniasis (75), or by promoting oedema to retain the virus at the inoculation site, as seen with *Aedes aegypti* mosquito bites (34). Since we only examined the influence of spot feeding by *S. calcitrans* it is not clear if the acceleration in disease development is specific to this arthropod species or more broadly applicable to other species known to transmit LSDV such as mosquitoes. In fact, the mechanism may not be specific to arthropods at all but could be due to any trauma, such as scarification which has been shown to influence the immune response to the poxvirus vaccinia virus (76).

Understanding the impact of *S. calcitrans* saliva and spot feeding on the kinetics of LSD provides an insight into the mechanical transmission of poxviruses. By studying the influence of *S. calcitrans* on the pathogenesis of a bovine disease in its natural host species, this work has revealed that the mechanical transmission of pathogens by arthropod vectors is more sophisticated and intricate than previously thought, and worthy of deeper investigation.

## Materials and Methods

### Viruses

LSDV was propagated on MDBK cells (ATCC code CCL-22) in high-glucose Dulbecco’s Modified Eagle’s medium (Life Technologies No. 41965) supplemented with 2.5% fetal bovine serum (Antibody Production Services Ltd), and 1% penicillin-streptomycin (Life Technologies No. 15140122) at 37°C in a 5% CO_2_ atmosphere. LSDV was semi purified through a 36% sucrose cushion prior to titration. Infectious virus titre expressed as number of plaque or foci forming units/millilitre (PFU/mL), was determined by titrating virus on MDBK cells as previously described (77).

### Isolation of primary bovine fibroblasts

Bovine primary fibroblasts (BPF) from the distal ear pinna of skin were isolated and expanded as previously described (77). Briefly, the dermis was dissociated from a section of skin before pieces were then placed in a tissue culture plate in growth medium overnight at 37°C, in 5% CO_2_ atmosphere. More media was then added and the skin pieces removed two days later. Adherent fibroblast colonies were then seeded into cell culture vessels and grown at 37°C, 5% CO_2_.

### Infection of fibroblasts

Fibroblasts were grown to 90% confluency in 6-well tissue culture plates in BPF media. The cells were infected with LSDV at a high MOI (5) or mock infected for 1h at 37°C in 5% CO_2_. The inoculum was then removed, the cells washed twice with media, then incubated at 37°C in 5% CO_2_. Cellular and supernatant fractions were collected together at timepoints up to 7 days post infection (dpi) and infectious virus titres determined by plaque assay on MDBK cells.

### Saliva collection

The feeding reservoirs of the Hemotek™ membrane feeding system were prepared by securing an unstretched layer of Parafilm™ with a rubber ring, sealing the inner chamber before the reservoir was filled with endotoxin free water (Fisher Scientific). A Durapore™ PVDF 0.2 μm membrane (Millipore) was dipped briefly in 1mM ATP/endotoxin free water solution before being laid over the parafilm. Two thinly stretched layers of Parafilm™ were laid on top of the PVDF to allow for penetration by the proboscis but minimise contamination of the membrane by non-salivary proteins during feeding (Figure 1). Excess Parafilm™ was trimmed away from the completed reservoir insert before it was screwed into the main Hemotek™ apparatus and allowed to warm to 37°C. Laboratory-reared male and female *S. calcitrans* at an average of 14.5 days post-eclosion (range: 10-19 days) were starved the day before feeding, then placed into 53mm diameter plastic pots closed with mesh netting and placed onto the membrane feeding system (MFS) for approximately 15 min per pot. After the MFS was fed on 3-4 times by a pot of flies the Durapore™ membrane was placed into PBS and stored at 4°C.

### Saliva concentration

Once saliva collection from the flies was completed, the membranes were cut into four irregular pieces and washed in PBS at 4°C overnight under constant rotation. The eluted proteins were then sterilised by passing the eluate through a 0.2 μm filter (Merck Millipore), then concentrated using a 3kDa Vivaspin 20 centrifugal concentrator column (Sartorius) to give a final volume of approximately 1000 μL. The protein concentration in the saliva was calculated using the Pierce™ BCA Protein Assay Kit (ThermoFisher Scientific).

### SDS-PAGE and silver stain/Coomassie

Whole cell lysates were prepared by reduction and denaturation with dithiothreitol (DTT) and heating for 5 mins at 95°C. 15% SDS-PAGE gels were then used to separate *S. calcitrans* salivary proteins by size with proteins visualised by Coomassie brilliant blue dye or silver staining (Pierce™ Silver Stain Kit, ThermoFisher Scientific) according to manufacturer’s instructions.

### Proteomic analysis of *S. calcitrans* secreted saliva

#### Sample digestion

In-gel digestion was essentially as described by Shevchenko et al. (78). Pre-excised and stained bands were cut into smaller pieces (approx. 1mm^3^). Gel pieces were then de-stained and in gel digestion performed as described below.

Slices were de-stained with 25mM ammonium bicarbonate/ 50 % acetonitrile (v/v). Proteins were reduced for 10 minutes at 60°C with 10 mM dithiothreitol (Sigma) in 25 mM ammonium bicarbonate and then alkylated with 55 mM iodoacetamide (Sigma) in 50 mM ammonium bicarbonate for 30 minutes in the dark at room temperature. Gel pieces were washed for 15 minutes in 50 mM ammonium bicarbonate and then dehydrated with 100% acetonitrile. Acetonitrile was removed and the gel plugs rehydrated with 0.01 µg/µL proteomic grade trypsin (Thermo Fisher Scientific) in 25 mM ammonium bicarbonate. Digestion was performed overnight at 37°C. Peptides were extracted with 50% (v/v) acetonitrile, 0.1% trifluoroacetic acid (TFA) (v/v) and the extracts were reduced to dryness using a centrifugal vacuum concentrator (Eppendorf) and re-suspended in 3 % (v/v) methanol, 0.1 % (v/v) TFA for analysis by MS.

#### Mass spectrometry

LC-MS/MS analysis was similar to that described by Aljabr et al. (79). Peptides were analysed by on-line nanoflow LC using the Ultimate 3000 nano system (Dionex/Thermo Fisher Scientific). Samples were loaded onto a trap column (Acclaim PepMap 100, 2 cm × 75 μm inner diameter, C18, 3 μm, 100 Å) at 9 μL /min with an aqueous solution containing 0.1 % (v/v) TFA and 2% (v/v) acetonitrile. After 3 min, the trap column was set in-line an analytical column (Easy-Spray PepMap® RSLC 50 cm × 75 μm inner diameter, C18, 2 μm, 100 Å) fused to a silica nano-electrospray emitter (Dionex). The column was operated at a constant temperature of 35°C and the LC system coupled to a Q-Exactive mass spectrometer (Thermo Fisher Scientific). Chromatography was performed with a buffer system consisting of 0.1 % formic acid (buffer A) and 80 % acetonitrile in 0.1 % formic acid (buffer B). The peptides were separated by a linear gradient of 3.8 – 50 % buffer B over 30 minutes at a flow rate of 300 nl/min. The Q-Exactive was operated in data-dependent mode with survey scans acquired at a resolution of 70,000 at m/z 200. Scan range was 300 to 2000m/z. Up to the top 10 most abundant isotope patterns with charge states +2 to +5 from the survey scan were selected with an isolation window of 2.0Th and fragmented by higher energy collisional dissociation with normalized collision energies of 30. The maximum ion injection times for the survey scan and the MS/MS scans were 250 and 50ms, respectively, and the ion target value was set to 1E6 for survey scans and 1E4 for the MS/MS scans. MS/MS events were acquired at a resolution of 17,500. Repetitive sequencing of peptides was minimized through dynamic exclusion of the sequenced peptides for 20s.

#### Protein Identification

Spectral data were analysed using the PEAKS studio X+ software (Bioinformatics Solutions Inc., Waterloo, ON, Canada (80)). Tandem MS data were searched against the *Stomoxys calcitrans* protein database (UniProt) and a contaminant database (cRAP, GPMDB, 2012). Search parameters were as follows; precursor mass tolerance set to 15 ppm and fragment mass tolerance set to 0.02 Da. Trypsin, was set as the protease. Two missed cleavage events were permitted. Carbamidomethylation (C) was set as a fixed modification. Oxidation (M) and acetylation (N-term) were set as variable modifications. The search was specific. The false discovery rate was set at 1%.

#### Sialome Functional Enrichment Analysis

Functional enrichment analysis of proteins represented in at least two out of three saliva collections was conducted using the g:GOSt algorithm available on the g:Profiler web server (81). UniProt accession numbers in a ranked list as shown in Table 1 were queried against the *S. calcitrans* Ensemble genome version GCA_001015335.1/SCIU1.6 (82) integrated in g:GOSt. Where alternative accession numbers for the same protein existed, these were removed prior to analysis. The g:SCS algorithm in g:GOSt was employed for computing multiple testing correction for p-values gained from the GO and pathway enrichment analysis with an experiment-wide p-value of 0.05 (confidence interval level of 95%).

##### Animal study

The animal study was performed in the high-containment animal facilities at The Pirbright Institute under project license P2137C5BC from the UK Home Office according to the Animals (Scientific Procedures) Act of 1986, and was approved by The Pirbright Institute Animal Welfare and Ethical Review Board. Thirty male Holstein-Friesian calves 4-6 months old (weight range 133-167kg) were included in the study. The study was carried out as two replicates each of 15 animals. In each replicate animals were stratified according to weight then randomly assigned to the three treatment groups. The animals were sourced from a commercial high health herd and confirmed as negative for bovine viral diarrhoea virus (BVDV) via PCR prior to study commencement. The animals were housed in 3 rooms (22m^2^) in a high-containment (SAPO4) animal facility at the Pirbright Institute. Bedding material was provided (https://www.mayofarmsystems.co.uk/mayo-mattress-stable-mat/), light/dark cycle was 12:12 h, temperature was held between 10°C to 24°C, and humidity 40% to 70%. Animals were fed concentrated rations twice daily and given ad lib access to hay and water. Environmental enrichment was provided, including rubber toys and a hollow ball stuffed with hay. Calves were treated for three co-morbidities during the study (ringworm, bronchopneumonia, and orchitis / epididymitis secondary to castration) as detailed in the supplementary data.

Calves in group 1 were inoculated with 1×10^6^ PFU/mL LSDV intradermally. The inoculum was delivered as ten 100 µl “microdoses” into an 8cm diameter region of skin on the left and right craniodorsal flank, just caudal to the scapula. Calves in group 2 were challenged with 1×10^6^ PFU/mL LSDV that had been mixed with 10μg *S. calcitrans* salivary proteins. Inoculum was delivered as in group 1. Calves in group 3 had two pots, each pot containing approximately 25 *S. calcitrans* flies, placed sequentially on the skin of the left and right craniodorsal flank for 5 min each pot. Each animal was then challenged with 1×10^6^ PFU/mL LSDV with the inoculum delivered in microdoses (as described for groups 1 and 2), into the site where the flies had fed.

Following inoculation each animal was clinically examined regularly including heart rate, temperature, respiratory rate, swelling at inoculation site, number of nodules (and their locations), anorexia, lameness, behavioural changes (including isolation, lethargy, abnormal posture), and lymphadenopathy. Venous blood samples and skin microbiopsies were collected at regular intervals up to 28 dpi.

##### Clinical scoring system

A scoring system for experimental LSD developed by Carn and Kitching (83) was modified to include the full range of LSD clinical signs seen after microdose inoculation. The scoring system included cutaneous nodules, enlargement of lymph nodes, temperature, unusual behaviour, anorexia, oedema and lameness, ocular/nasal discharge and salivation. These criteria were then weighted to account for severity of some clinical signs (Table 3).

##### PCR of blood

Nucleic acid extraction and PCR was conducted by the Non-Vesicular Reference Laboratory at The Pirbright Institute. DNA was extracted using the MagMAX™ CORE Nucleic Acid Purification Kit (Applied Biosystems) on a Kingfisher flex automated extraction robot (ThermoFisher Scientific). The starting sample volume was 200 µL, and the DNA was eluted into 90 µL elution buffer in the final step. The real-time PCR assay for CPPV (Bowden et al., 2008) targeted an 89 bp region within the LSDV074 ORF. Each assay used a dual labelled fluorogenic (TaqMan®) probe that allowed the assay to be performed in a closed-tube format minimising the potential for cross-contamination of post-PCR products.

##### Titration of blood

PCR positive blood samples were serially diluted in calcium-free DMEM, before infectious virus titre (number of plaque-forming units/millilitre, PFU/mL) was determined by plaque assay on MDBK cells (77).

##### Skin biopsy PCR

Experiment was carried out according to the protocol used by Sanz-Bernardo et al. (84). Viraemia was analysed in duplicate, and results presented as mean copies per 2mm biopsy.

##### Flow cytometry

Antibody cocktails were prepared with 0.5% bovine serum albumin (BSA)/PBS as diluent. 100 µL heparinised whole blood from each animal was aliquoted in triplicate into empty Round-bottom polystyrene 5ml (Falcon 5ml) tubes and mixed with 30 µL antibody cocktail, made of CD3 (MM1a) conjugated with AF488 (Alexa Fluor™ 488 Antibody Labeling Kit), CD14 (CC-G33, BioRad) conjugated with AF647 (Alexa Fluor™ 647 Antibody Labeling Kit), CD40 (IL-A158, Immunological Toolbox) conjugated with PE (Lightning Link, Novus Biologics), and γδ TCR (GB21a, Washington State University) conjugated with PB (Pacific Blue Antibody Labeling Kit). Blood from CV1 and CV6 cattle were used as single colour (SC), fluorescence minus one (FMO) and unlabelled controls. The blood samples and antibodies were incubated for 30 min at RT in the dark. 2 mL eBioscience™ Fix/Lyse Solution (10×) diluted 1:10 in sterile water was added to each tube to lyse the red blood cells. The tubes were incubated at RT for 30 min in the dark. 100µl of vortexed Invitrogen™ 123count eBeads™ (Fisher Scientific) were added to each tube. Samples were read on a 4 Laser LSR Fortessa (405nm, 488nm, 561nm and 640 nm). AF488 signal was detected with BP530/30, after excitation with the 488nm laser. AF647 signal was detected with BP670/14, after excitation with the 640nm laser. PE signal was detected with BP582/15, after excitation with the 561nm laser. PB signal was detected with BP450/50, after excitation with the 405nm laser. Compensation was run on single colour controls. Cells were gated according to **Figure 11**, and data was analysed in FCS Express 7 (De Novo software).

**Figure 11.**
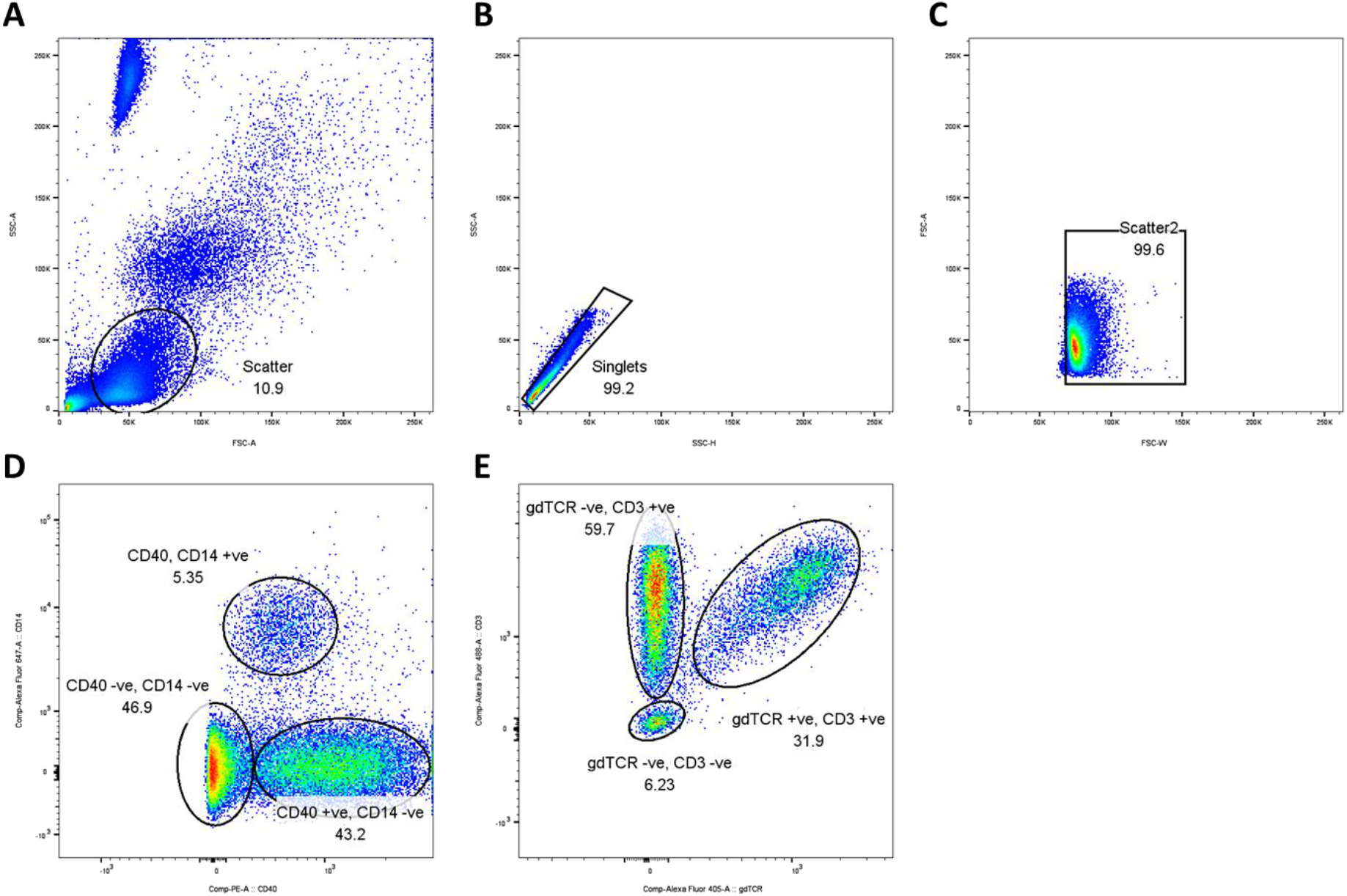
The whole blood flow cytometry panel gating strategy. Cells were plotted on a forward vs side scatter plot (A) to isolate the mononuclear cell population, then gated to exclude doublets via a side scatter area vs height plot (B). Further debris was excluded by gating on a forward scatter area vs width plot (C). Monocyte (CD14 positive, CD40 negative) and B cell (CD14 negative, CD40 positive) populations were selected by plotting markers for CD14 vs CD40 (D). The CD40/CD14 negative population was plotted on CD3 vs gamma delta T cell receptor (γδ TCR) cell marker plot, to isolate the T cell population (CD3 positive, γδ TCR negative) and gamma delta T cells (CD3 positive, γδ TCR positive) (E).

##### Fluorescent Virus Neutralisation Test

The neutralisation titre was determined as described previously (71). Briefly, MDBK cells were seeded (3×10^5^/mL) in 96 well plates. Sera were heat-inactivated at 56°C for 30 min then diluted 1:10 in cell culture medium (DMEM, 2.5% heat-inactivated FBS, 100 units/mL penicillin, and 100µg/mL streptomycin). Two-fold serial dilutions from 1:10 to 1:1280 were prepared. Recombinant EGFP-095-LSDV Neethling virus (7×10^2^ pfu/mL) was added to each dilution and incubated for 1h at 37°C in a 5% CO_2_ atmosphere. Samples were added to the MDBK cells and incubated at 37°C in a 5% CO2 atmosphere for 5 days. To interpret the results, fluorescent foci were counted using a fluorescent UV light microscope (Olympus CKX53). The complete neutralising antibody titre was calculated as the highest dilution at which no foci were identified. All samples and controls were tested in duplicate.

### Statistical methods

#### Clinical scoring

Mixed effects models were conducted to determine statistical significance between the treatment groups and clinical outcomes. p-values < 0.05 were considered to be statistically significant.

#### Nodule number

Mixed effects models were conducted to determine statistical significance between the treatment groups and clinical outcomes. p-values < 0.05 were considered to be statistically significant.

#### FVNT

Because of non-normality of errors and non-equal variances, the median log FVNT were compared amongst calves infected by each method (Group 1, Group 2 or Group 3) at each timepoint. If the Kruskal-Wallis test was significant (P<0.05), pairwise Wilcoxon rank-sum tests (two-sided, unpaired) were used to compare responses between groups.

#### % CD14+ cells

Because of non-normality of errors and non-equal variances, the median percentage CD14+ cells were compared amongst clinical and non-clinical calves (regardless of infection method) at each timepoint using a Wilcoxon rank-sum test (two-sided, unpaired).

#### C_T_ values in blood

Because of non-normality of errors and non-equal variances, median CT values in blood were compared amongst groups at each timepoint. Calves were divided into six groups based on the infection method (Group 1, Group 2 or Group 3) and clinical status (clinical or non-clinical). If the Kruskal-Wallis test was significant (P<0.05), pairwise Wilcoxon rank-sum tests (two-sided, unpaired) were used to compare responses between groups, specifically: (i) clinical and non-clinical cattle infected by the same method; (ii) clinical cattle infected by different methods; and (iii) non-clinical cattle infected by different methods.

A sample with no C_T_ value was given an arbitrary value of 42 for the purposes of analysis. In addition, samples were tested in duplicate. The mean of the duplicate values for each sample was used in the analysis, unless one result produced a C_T_ value and the other did not, in which case only the (positive) C_T_ value was used.

The analyses of clinical scores and nodule numbers were implemented in GraphPad Prism 8, while the analyses of FVNT, % CD14^+^ and C_T_ values were implemented in R (version 4.1.3) (85).

## Acknowledgements

The authors are grateful to the Animal Services and the Non-Vesicular Reference Laboratory staff at The Pirbright Institute for their invaluable assistance.

## Funding

This project received funding and support from BBSRC responsive mode project BB/T005173/1, and BBSRC strategic funding to the Pirbright Institute BBS/E/I/00007030, BBS/E/I/00007031, BBS/E/I/00007033, BBS/E/I/00007036, BBS/E/I/00007037, BBS/E/I/00007039. The project received funding from MSD Animal Health, and from the European Union’s Horizon2020 research and innovation programme under grant agreement No 773701.

## Author contributions

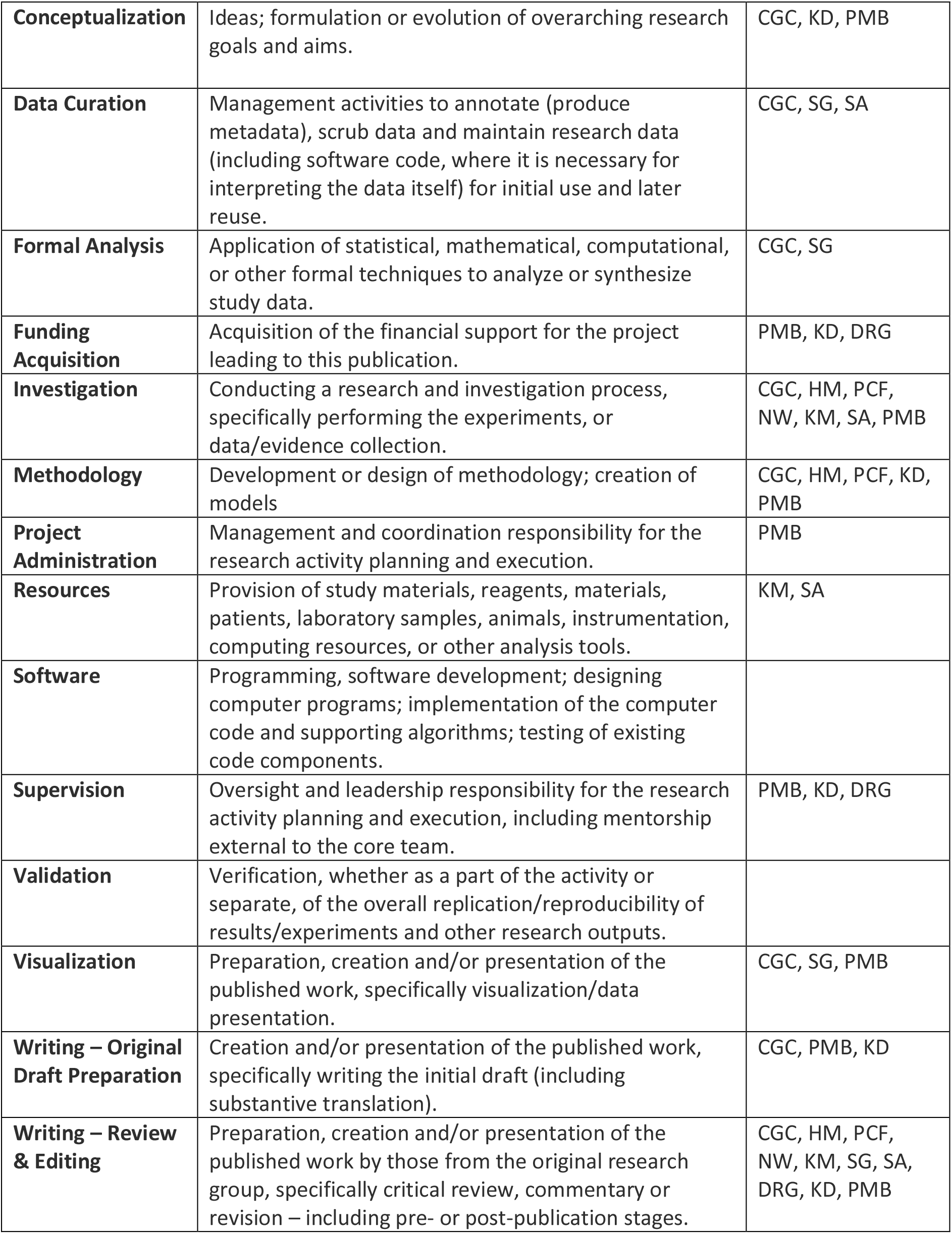

## Notes

### Competing Interest Statement

The authors have declared no competing interest.

